# Sulforaphane targets *Leishmania* promastigotes to inhibit parasite motility and alters host autophagy to suppress parasite replication in macrophages

**DOI:** 10.64898/2026.09.04.749349

**Authors:** Elenora Kelley, Emmet Thompson, Ahliya Harding, Katie Torrey, Kai Gurnoe Brantley, Chunyu Lin, Anders Bach, Kenneth A. Miller, David J. Blake

## Abstract

2.

Visceral leishmaniasis (VL) is a neglected tropical disease caused by the protozoan parasites *Leishmania infantum* (LI) and *Leishmania donovani* (LD). VL disproportionately affects populations that live in extreme poverty and those affected by climate change. Treatment options are limited, fiscally prohibitive, and toxic to some that receive treatment. In cases where VL remains untreated, death is inevitable. Although disease prevalence is increasing in affluent countries with robust research capabilities, there remains little economic incentive to search for new, more accessible non-toxic treatments, emphasizing the neglected tropical aspect of this disease. Our previous results indicate that sulforaphane (SFN) inhibits VL promastigote and amastigote growth through an unknown mechanism. The current study extends the anti-parasitic effects of SFN establishing SFN’s efficacy against multiple *Leishmania* parasites including *L. tropica, L. mexicana, L. panamensis* and *L. amazonesis*. Moreover, SFN was an effective therapy post-infection as stably infected macrophages treated with SFN (10 µM for 48 hours) had a significant reduction in the number of infected cells and the number of amastigotes per cell against *Leishmania* species that cause both VL and cutaneous leishmaniasis (CL). Moreover, SFN led to a reduction of motility within 6 hours of treatment indicating a direct interaction with the parasite. Alkynyl SFN (A-SFN) was subsequently synthesized to determine the specific binding target of SFN. Interestingly, A-SFN was more effective than SFN in decreasing parasite viability and inhibiting motility against VL and CL parasite species at equivalent concentrations. Immunofluorescence analysis utilizing click chemistry indicated A-SFN binds to cytosolic promastigote components and colocalized only minimally with KMP-11. The binding of SFN to KMP-11 could not be confirmed *in vitro.* In terms of intracellular host effects, SFN treatment increased the induction of nonselective macroautophagy by increased lysosomal acidification as quantified by DALGreen and FITC-dextran staining in infected macrophages. The induction of autophagy occurred simultaneously with the upregulation of expression of cytoprotective, antioxidant and autophagy-related genes via the NRF2 transcription factor. These data provide robust evidence that SFN is an excellent lead compound against *Leishmania* that initiates multiple innate immune pathways that enable enhanced parasite killing. Future studies will focus on the *in vivo* efficacy of SFN against CL.

**Graphical abstract:** 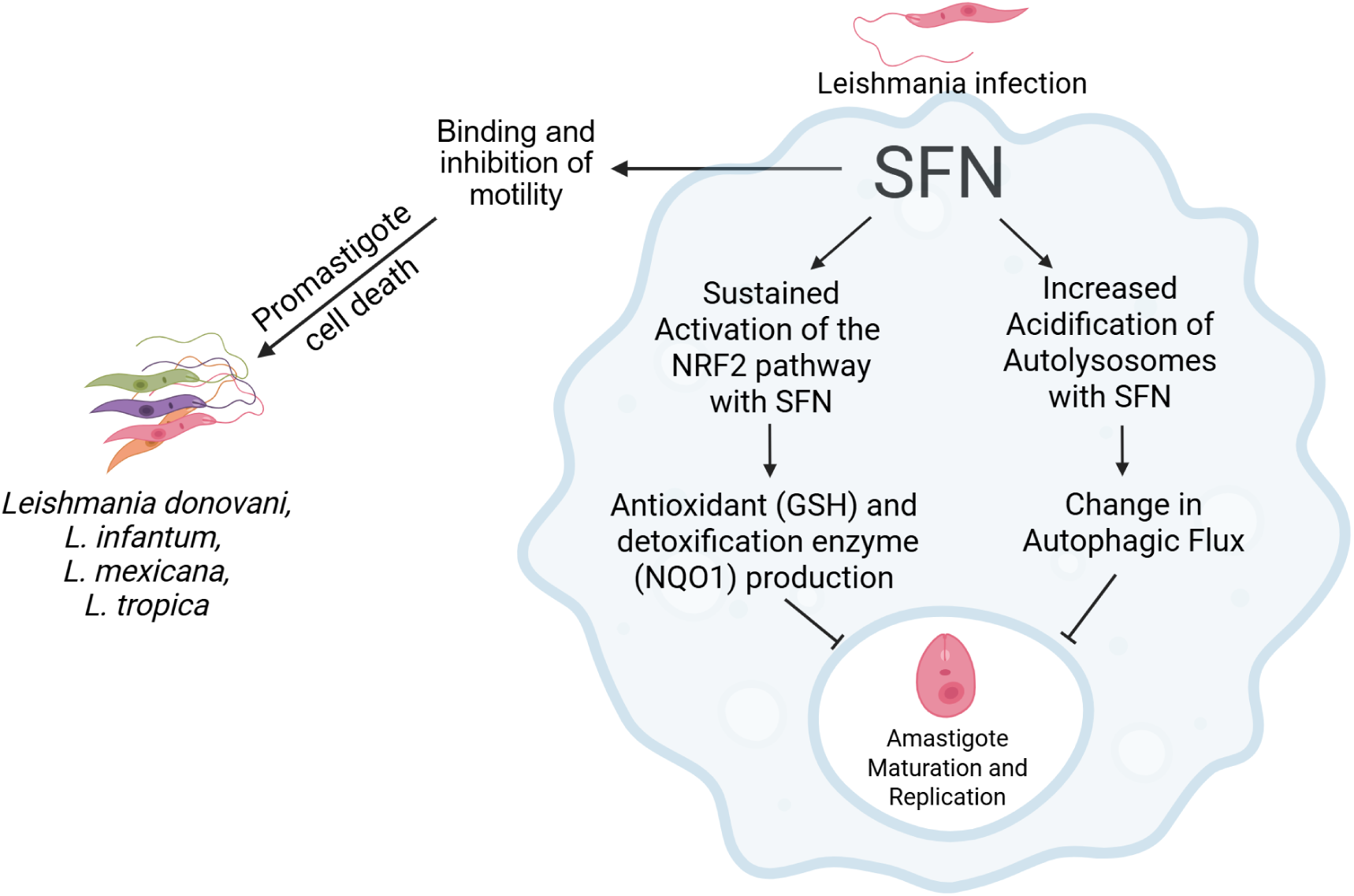

**KEY FINDINGS:**

- Sulforaphane (SFN) decreased promastigote viability from six *Leishmania* species
- SFN is an effective therapy that decreased the number of intracellular VL and CL amastigotes in macrophages post-infection
- SFN inhibited promastigote motility within 6 hours
- SFN increased NRF2-dependent pathway expression and lysosomal acidification in cells infected by *Leishmania*

## 4. INTRODUCTION

Leishmaniasis is a vector-borne neglected tropical disease (NTD) transmitted by sandflies through the infection of over 20 different *Leishmania* species. Infection occurs in approximately 1 million people each year leading to more than 50,000 deaths annually, making leishmaniasis the second largest parasitic killer in the world after malaria [1–3]. Visceral leishmaniasis (VL) is the deadliest clinical form of the disease caused by *Leishmania donovani* (LD) and *L. infantum* (LI) resulting in death in 95% of untreated cases [4]. Cutaneous leishmaniasis (CL) is the majority of infections worldwide causing skin infections and disfiguring lesions of the face and mucosal membranes due to *L. mexicana, L. tropica, L. panamensis, or L. amazonensis* infections (Pareyn et al., 2025). No universal treatment exists, thus, antileishmanial therapies vary by region and are dependent on the parasite species.

The lack of alternative treatments is a consequence of limited investment for NTDs that primarily affect low-income countries and exacerbates effective treatment options [6]. Moreover, current therapies have several drawbacks, including reduced efficacy, toxicity, and need for hospitalization that increases treatment costs significantly. Unfortunately, the industrial pipeline is very limited, and no therapeutic breakthroughs are expected [7]. The lack of cost-effective therapies juxtaposed with an increasing number of endemic countries infected with *Leishmania,* including the USA beginning in 2023, highlights the need for new compounds to be investigated [8–12]. Ideally, any new therapy would be effective across all *Leishmania* species, have limited toxicity, be an accessible treatment to the communities without the need for hospitalization, and in the best case have multi-target effects that promote host innate immunity to eliminate the infection, thus increasing the likelihood of *in vivo* efficacy.

Kinetoplastid Membrane Protein-11 (KMP-11) is a major surface protein expressed in many different kinetoplastids including *Leishmania* promastigotes and amastigotes [13] as well as in *Trypanosoma brucei* where KMP-11 is essential in cell division [14]. The conservation of KMP-11 and its wide distribution in kinetoplastids suggests an essential role for the parasite [15]. In *Leishmania*, KMP-11 is found at the cell surface, at the flagellar pocket and within intracellular vesicles [16]. KMP-11 is a highly conserved protein among *Leishmania* species (Ramírez et al., 1998) although the number of exons is variable among species. Immunologically KMP-11 acts as a virulence factor that promotes IL-10 secretion to downregulate IFN-γ synthesis and induces a T_H1_ cytokine response that promotes parasite elimination [18]. The biological purpose of KMP-11 has been linked to parasite motility, adherence and infection in macrophages (MΦs), induction of T cell proliferation, and cytoskeleton regulation [19–21]. Recently, KMP-11 was shown to facilitate cholesterol transport to the parasite while remaining on the parasite surface during infection and subsequent invasion [21]. The importance of KMP-11 during infection was supported by studies showing that KMP-11 depleted LD parasites fail to attach and invade host MΦs [21].

Previous *in silico* studies indicate that KMP-11 can be bound by natural products such as sulforaphane (SFN). SFN is a naturally occurring isothiocyanate (ITC) found in cruciferous vegetables [22]. SFN’s *in vitro* effectiveness was confirmed by our laboratory and identified antileishmanial effects of SFN against both LD and LI promastigotes and intracellular amastigotes [23]. However, SFN is a complex phytochemical that can activate broad cellular pathways compared to more specific NRF2 activators [24]. We initially hypothesized that SFN and its reactive isothiocyanate group directly binds to KMP-11 to inhibit motility and decrease macrophage (MΦ) infection.

Our results indicate that SFN is an effective therapy against multiple *Leishmania* species as promastigotes or amastigotes, and inhibits motility within 6 hours. Synthesizing A-SFN and utilizing click chemistry identified binding of SFN to a cytoplasmic component within Leishmania promastigotes. However, the binding of SFN to KMP-11 was not supported. SFN also induces beneficial host responses, such as increased lysosomal acidification, which may contribute to SFN’s multi-target antiparasitic mechanisms. These results provide adequate mechanistic understanding of SFN to investigate its effects *in vivo*.

## 5. MATERIALS AND METHODS

### General Experimental Procedures

All general chemicals were purchased from Sigma-Aldrich (St. Louis, MO). RPMI 1640 media, M199 media, fetal bovine serum, antibiotics (penicillin/streptomycin) solution, propidium iodine, FITC-dextran [46945], KMP-11 [L-157] and rabbit anti-mouse Alexa Fluor 488 and AF594, Pierce NeutraAviden UltraLink Resin, and Prolong Gold mounting media were purchased from Thermo Fisher Scientific (Waltham, MA). CellTiterBlue reagent was purchased from Promega (Madison, WI). R-Sulforaphane was purchase from LKT laboratories (St. Paul, MN). Protein-protein inhibitors (compounds UCAB#985 and UCAB#1124) were synthesized as previously described in the Bach lab [24]. 4% PFA was purchased from Biotium (Fremont, CA). His+SUMO combination tagged recombinant KMP-11 proteins from *Leishmania donovani* [VAng-Wyb7425] and *infantum* [VAng-Wyq9292] were obtained from Creative Biolabs Vaccine (Shirely, NY). Sequences for both KMP-11 proteins are included in Figure 5. Biotin Azide Plus was obtained from Vector Laboratories (Newark, CA). Coverslips (1.5H) were obtained from Carl Roth (Germany). DALGreen was purchased from Dojindo Laboratories (Rockville, MD).

### Leishmania and Human Macrophage Cell Culture

*Leishmania donovani* (LD) strain MHOM/IN/DD8 (Laveran and Mesnil) Ross, *L. infantum* strain MHOM/TN/80/IPT-1 Nicolle, *L. mexicana* strain MHOM/BZ/82/BEL21 (Biagi) Garnham, *Leishmania mexicana* MNYC/BZ/62/M379 (+*luc*), *L. tropica/major* strain (Yakimoff and Schokhor), *Leishmania amazonensis*, Strain LV78 (+bla), NR-49247, NIAID, NIH: *Leishmania panamensis*, Strain PSC-1 (MHOM/PA/94/PSC 1), NR-50162, and the THP-1 human monocytic cell line were purchased from ATCC or the BEI Resources NIAID (Manassas, VA). Leishmania parasites were cultured in M199 media supplemented with 10% FBS, antibiotics and heme in M199 media as previously described [25]. *Leishmania* cultures were maintained at 27°C, checked weekly for purity and subcultured as suggested by the manufacturer usually once every five days when the number of metacyclic promastigotes were at the highest concentrations. All parasite cultures used for all infections were passaged less than 7 times. THP-1 cells were maintained in RPMI supplemented with 10% FBS and antibiotics at 37°C in 5% CO_2_. THP-1 cells were subcultured every three days. All THP-1 cells used for infections were passaged less than 10 times.

### Promastigote Viability Assay

Promastigotes in the infective metacyclic stage (long cylindrical forms, ∼5 day old cultures for all cultures except LM ∼3 days old), approximately 1 million cells per well, were cultured in the presence of either 0.5 μM amphotericin B, 10 or 50 μM SFN for 48 h (n = 5–10 technical replicates). CellTiterBlue was added and incubated 3 h at 37 ◦C. Fluorescence was measured at (560Ex/590Em) in an Infinite 200 PRO series plate reader (Tecan Systems Inc., San Jose, CA). Amphotericin B was used as the positive control and M199 media was used as the negative control. Percent viability was calculated for each well after background subtraction, each well was compared to the average control using the following equation; percent viability = (590Em _sample_/590Em _mean negative control_) * 100. Each data point represents a mean viability ± SD and was normalized to the mean value of the negative control corresponding to identical species and experiment. Experiments were repeated at least three times (≥3 biological replicates).

### Assessment of Antileishmanial Activity of SFN Post-Infection

THP-1 cells were differentiated into macrophage-like cells for 48 hours with phorbol 12-myristate 13-acetate (PMA) at a final concentration 25 ng/ml. 5-day old metacyclic LD promastigotes were used to infect adherent macrophages at either a 20:1 or 10:1 ratio (promastigotes to human macrophages) for 5 h in RPMI media at 37°C. Cells were washed twice with PBS to remove unattached promastigotes and then incubated for an additional 18 hours. Cells were then washed and treated with 10 μM SFN for 48 h (n = 7-10 technical replicates). Approximately 80% of macrophages are infected after 48 hours with a 20:1 ratio for 5 hours (unpublished results). Alternatively, cells were infected with a 20:1 ratio for 24 hours continuously, then washed and exposed to 10 μM SFN for an additional 48 h (n = 7 technical replicates). Cells were washed with PBS twice, fixed with 4% paraformaldehyde, and permeabilized. Nuclei were identified by staining with DAPI. All samples were blinded prior to image acquisition. All images were obtained with an Olympus BX60 microscope and CellSens software. Human macrophages (large nuclei) and amastigote (small punctate nuclei) were enumerated as blinded samples to quantify the number of amastigotes per 100 THP-1 cells. Experiments were repeated twice (2 biological replicates).

### Quantification of Promastigote Motility

Low magnification videomicrographs for motility analysis were captured using bright field illumination at a frame rate of 5 Hz, using a 10× NA 0.25 objective on a IX50 inverted microscope (Olympus Microsystems) and an DP73 camera (Olympus) with an exposure time of 1 ms as previously described [26]. Cell culture conditions were maintained in 50 ml conical tubes at 27°C. Each condition was transferred into 6-well plates (Fischer Scientific) and imaged after 3 and 6 hours (n = 25-120 parasites per time point). Motility from low magnification videomicrographs was quantified using KTSam2, a custom automated pipeline built on Meta’s Segment Anything Model 2 (SAM2). Briefly, individual videos were broken into single frames, and a mask was manually drawn on the first frame in Fiji (ImageJ). SAM2 utilizes the mask as a reference to track all parasites through all frames. Parasite center points were then extracted from each mask and linked into trajectories using Trackpy. Gaps up to 3 frames were allowed before ending the track. Tracks were built within a maximum connection distance of 15 pixels (3.2 μm). If no cell could be found within that range, then the track was terminated. In each track, the following was tracked: average speed and total distance travelled. A sample of the parental cell line killed with a final concentration of 1% paraformaldehyde was used as a reference for motion of completely paralyzed cells through sedimentation and Brownian motion alone. This background rate was subtracted out of all conditions within a single experiment. The cited method includes the use of PFA to immobilize parasites as a non-motile control, which was included in our method. In addition, a second positive control (200 mM H_2_O_2_) was utilized to confirm inhibition of motility. Experiments were repeated at least three times (≥ 3 biological replicates). Data are represented in violin plots that show the frequency of distribution of data. Medians and quartiles are indicated for each group.

### Synthesis of Alkynyl SFN (A-SFN)

Alkynyl SFN synthesis was followed as previously described [27,28].

### Synthesis of KEAP1 protein-protein interaction inhibitors

KEAP1 PPI inhibitors were created as previously described [24].

### Click reaction conditions and A-SFN localization via immunofluorescence

5-day old metacyclic promastigotes were treated with 100 µM A-SFN for 3 hours at 27 °C. Promastigotes were washed in PBS, fixed with 4% PFA at room temperature for 20 minutes, washed in PBS, attached by centrifugation onto poly-L-lysine-coated slides and permeabilized with 0.1% Triton in PBS as previously described [29]. For the Copper (Cu(I)-catalyzed Azide-Alkyne click) (CuAAC) chemistry reactions, crosslinked and permeabilized promastigotes were incubated with 5 µM AZDye488-Picolyl-Azide in a reaction buffer containing 2 mM CuSO_4_, 10 mM BTTAA and 100 mM Na-Ascorbate as suggested by the manufacturer (Jena Bioscience, Jena Germany). Reactions were incubated for 60 minutes at room temperature. The reaction was washed extensively and blocked in a solution containing 2% BSA/5% FBS for 1 hour at room temperature, then incubated with primary antibody (mouse anti-KMP-11 [L-157]) for one hour. Cells were washed in PBS three times and incubated with anti-mouse AF594-labeled secondary antibody. Samples were imaged with a BX60 microscope (Olympus Microsystems) and a DP73 camera (Olympus) or a ZIESS LCM 900 confocal microscope (UNM AIM Core).

### *In vitro* binding studies

Recombinant KMP-11 was incubated with A-SFN at a 250X molar excess for 4 hours at 4 °C as previously described [30]. After the initial incubation, click reactions were performed as described above utilizing 2 mM CuSO_4_, 10 mM BTTAA and 100 mM Na-Ascorbate and were incubated with 20 µM Biotin Azide Plus for 90 minutes at room temperature in the dark. After the click reaction reached completion, 50 µL NeutraAviden UltraLink beads were added overnight at 4°C. Precipitated complexes were washed twice in binding buffer and then resuspended in 2X Laemmli buffer. Precipitated complexes were resolved by standard SDS-PAGE conditions and stained with Coomassie.

### Image Analysis

Individual immunofluorescence images were collected on a ZIESS LCM 900 confocal microscope (UNM AIM Core) and colocalization of A-SFN with KMP-11 was assessed through ImageJ using the JACoP plug-in as previously described [31]. Briefly, green and red channels were spilt in Image J (Version 1.52a). Thresholds were set equally for both the green and red images (80). Finally, Mander’s overlap coefficient was calculated to determine colocalization. The M2 parameter quantified the proportion of green (A-SFN) with a signal in the red channel (KMP-11) over its total intensity.

### Quantification of autolysosomes and lysosomal acidification

Differentiated THP-1 cells were stained 1 µM with DALGreen in HBSS for 30 mins at 37°C, washed with PBS, and infected with LD promastigotes at a 20:1 infection ratio for 5 hours. Unattached promastigotes were removed after 5 hours through a second PBS wash. SFN was added to cultures post-infection. Cells were also exposed to 100 nM Bafilomycin A to inhibit v-ATPase and lysosomal fusion and served as the positive control to decrease DALGreen fluorescence associated with decreased numbers of autolysosomes. Cells were maintained for 18 hours post-infection. Alternatively, FITC-dextran 50 ug/ml) was added to cells after parasite infection and SFN treatment for three hours prior to quantification. Adherent cells were released with trypsin, which was neutralized with cold 10% FBS RPMI media, pelleted, washed and resuspended in HBSS as suggested by the manufacturer prior to analysis through flow cytometry. DALGreen fluorescence was quantified using the B1 channel (488 nM laser 525/50 nm filter) with a Miltenyi MACSQuant 10 (San Diego, CA). Cells were initially plotted and gated on a FSC vs. SSC plot, then single cells were identified through plotting the gated population on a FSC height vs FSC area plot. Only viable cells (PI negative) were quantified, and promastigotes were excluded from the analysis by including a trigger threshold of 25. Data represent either the mean or median green fluorescence (±SD) and the percentage of cells that were positive for green fluorescence. Experiments were repeated at least three times (≥3 biological replicates).

### Statistical analysis

Data are given as mean or median ± SD or mean ± SEM. Analyses were done using the software package GraphPad Prism 11 (GraphPad, San Diego, CA). One-way ANOVA was used to compare groups with one independent variable. Dunnett’s posttest was used to compare different treatments. Significance was noted at *P* < 0.05.

## 6. RESULTS

### SFN decreases promastigote and amastigote viability

The antileishmanial properties of SFN against LD and LI promastigotes as well as SFN’s ability to reduce LD amastigote levels has been previously reported (Leary et al., 2023) (Figure 1A and B). The antileishmanial effects of SFN can now extend against promastigote species that cause CL, as SFN significantly decreased the viability of *L. mexicana* (LM), *L. tropica* (LT)*, L. panamensis* (LP), and *L. amazonensis* (LA) at both 10 and 50 µM after 48 hours (Figure 1C-F). To explore whether NRF2 activation in *Leishmania* led to parasite cell death, two different noncovalent KEAP1-NRF2 protein-protein interaction (PPI) inhibitors (compounds UCAB#985 and UCAB#1124) were incubated with LD, LI, LT or LM promastigotes at 10 and 50 µM (Supplemental Figure 1). After 48 hours, no change in viability was observed for any *Leishmania* strain (data not shown) indicating an NRF2-independent pathway leads to parasite cell death.

**Figure 1:**
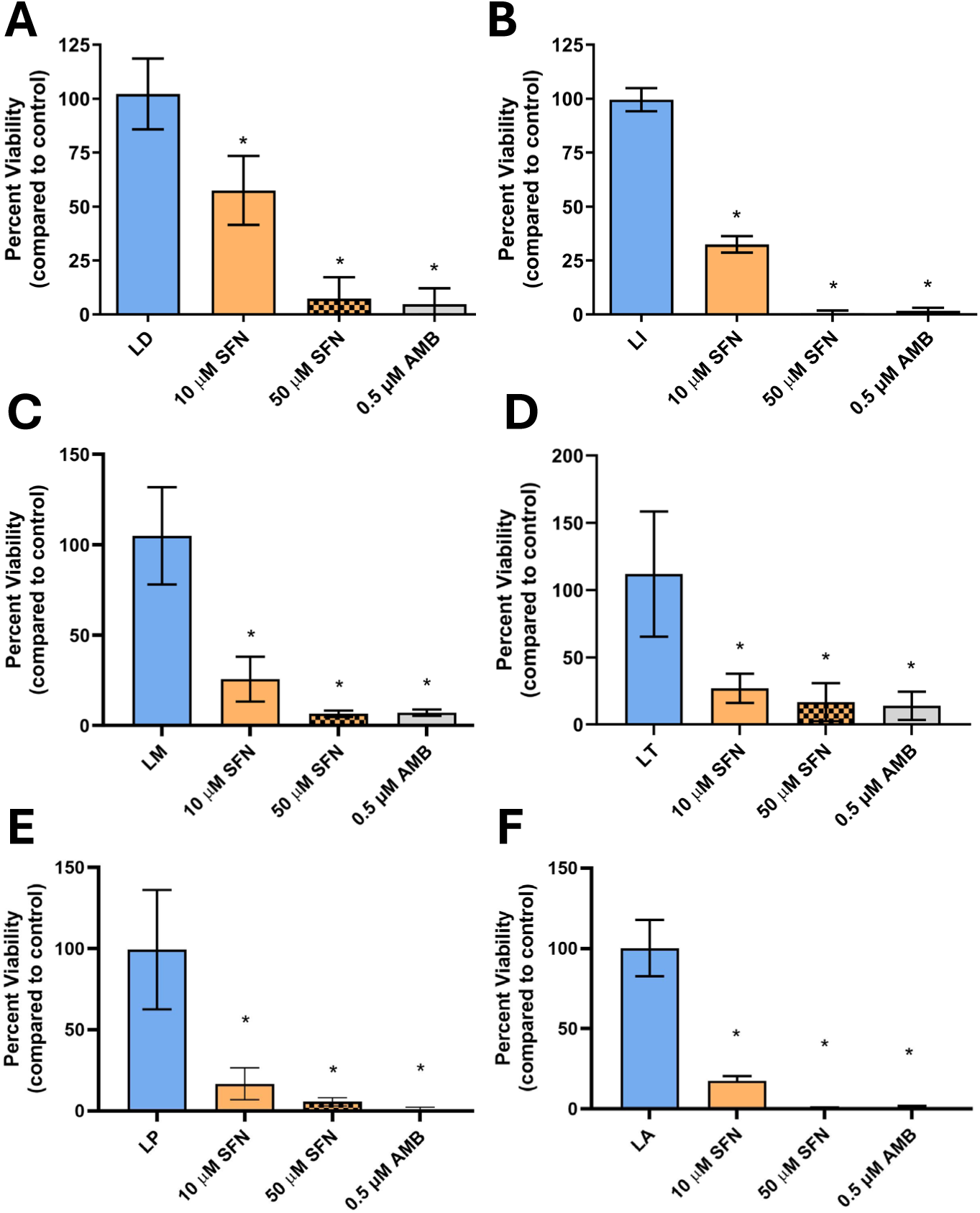
Sulforaphane is an effective anti-parasitic compound against six different *Leishmania* species. SFN decreases promastigote viability against *Leishmania donovani* and *L. infantum* **(A and B),** *L. mexicana* and *L. tropica* **(C and D)** and *L. panamensis* and *L. amazoneisis* **(E and F)**. Asterisks indicate a significant difference (*P* < 0.05) compared to control cells as determined through a one-way ANOVA and Dunnett’s post-hoc test. Data represent mean ± SD. Figures A and B have been modified from Leary *et al.* (2023). Figures C, D, E and F represent three biological replicates.

Because amastigotes are the parasite forms that persist in the infected host and the target of future treatments, MΦ’s were also stably infected with either LD or LI for 24 hours, then treated with 10 µM SFN for 48 hours. Infected cells treated with SFN had significantly lower amastigote numbers and infection levels compared to vehicle-treated controls (Figure 2A and B). SFN reduced amastigote levels by two-fold after 2 days of treatment, leading to a 30% reduction in infection levels. A single SFN dose did not lead to a significant reduction in the overall infection level or amastigote levels 5 days post-infection (p.i.). Therefore, multiple treatments of SFN may be needed over a 5-day period for significant parasite elimination. The anti-parasitic effects of SFN post-infection were confirmed in cells infected with *L. amazonensis*, which is a species that leads to cutaneous leishmaniasis (Supplemental Figures 2 and 3). These data indicate a direct anti-parasitic effect of SFN against the promastigote life stage of *Leishmania* and indicate intracellular efficacy post-infection as demonstrated by the elimination of intracellular amastigotes in stably infected MΦ’s (Figures 2A and B and Sup. Figures 2 and 3). Therefore, we interrogated the antileishmanial properties of SFN that are dependent upon direct parasite interactions and host-associated changes involving autophagic flux and the NRF2 pathway.

**Figure 2:**
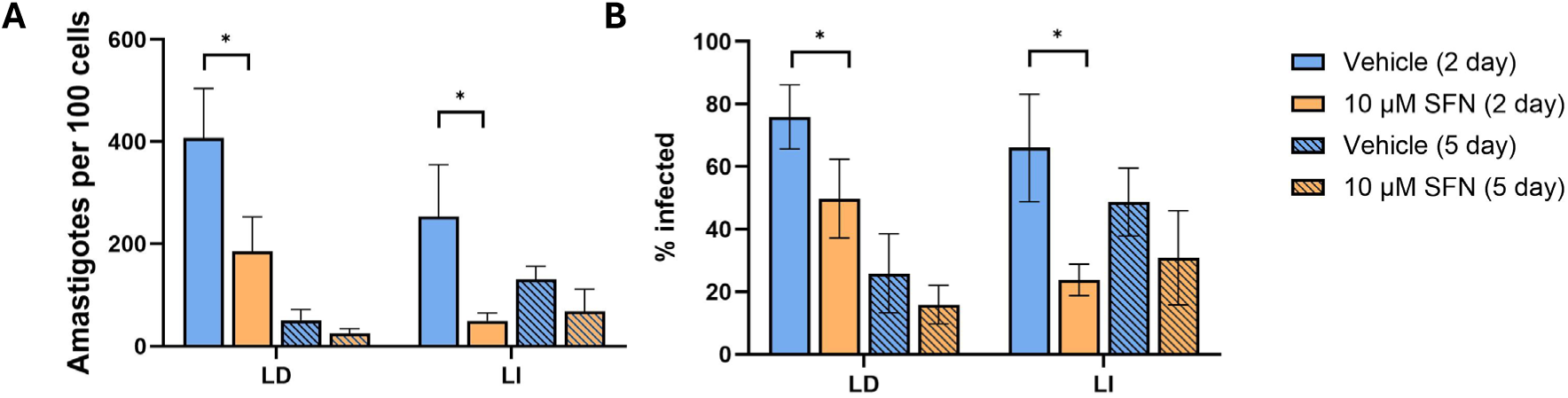
Sulforaphane is an effective therapeutic in macrophages stably infected with *Leishmania donovani* and *L. infantum*. Differentiated macrophages were infected with *L. donavani* (LD) and *L. infantum* (LI) for 24 hours. SFN treatment was given as a therapy 24-hour post-infection. **(A)** Parasite load, as quantified by the number of intracellular amastigotes per 100 macrophages, was significantly decreased after 2 days of SFN treatment. **(B)** The percentage of infected cells was quantified by enumerating the number of infected cells / total number of cells X 100. Asterisks indicate a significant difference (*P* < 0.05) compared to control cells as determined through a one-way ANOVA and Dunnett’s post-hoc test. Data represent mean ± SD and three biological replicates.

### SFN inhibits promastigote motility

To determine SFN’s effect on parasite motility, metacyclic LD promastigotes were treated with either 0, 10, 50 or 100 µM SFN at 27°C for three or six hours. Parasite velocities (µmeter/sec) and total distance migrated were quantified through a modified light microscopy protocol using video recordings after exposure [26]. No significant changes in motility were observed 3 and 6 hours after exposure to 10 µM SFN (Figure 3). However, both 50 and 100 µM SFN lead to significant reductions in parasite motility 3 and 6 hours after exposure. These reductions in velocity were accompanied by significant reductions in the total distances migrated by LD promastigotes after 3 and 6 hours of SFN at 50 and 100 µM. The inhibitory effects of SFN on motility were confirmed in a cutaneous leishmaniasis strain; *L. mexicana* (data not shown). The dose-dependent reduction in parasite velocity supports our hypothesis that SFN binds an essential *Leishmania* protein leading to an inhibition of motility.

**Figure 3:**
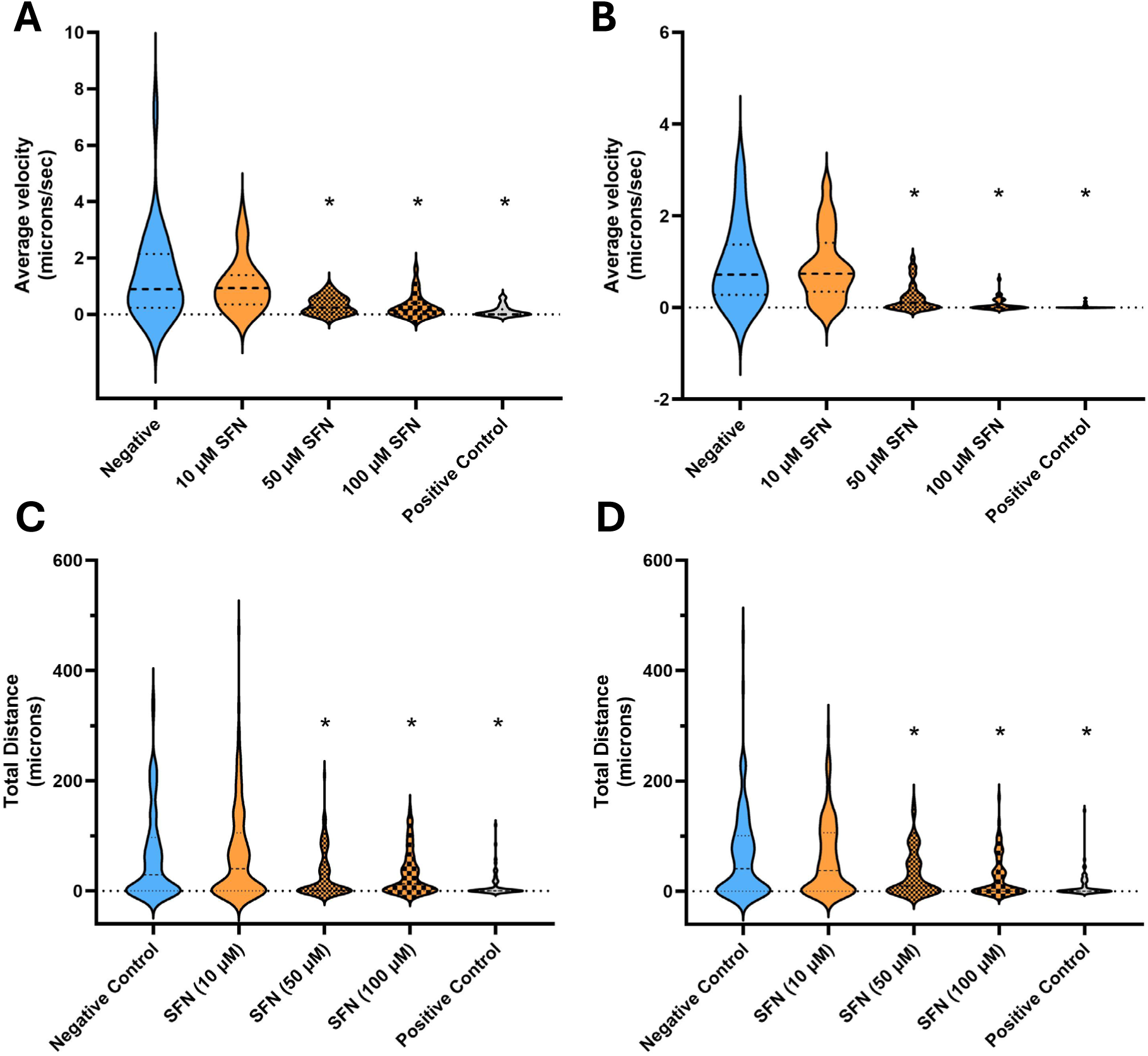
Sulforaphane inhibits *Leishmania* motility. LD promastigotes were exposed to SFN for 3 hours **(A and C)** or 6 hours **(B and D)** at 27 °C at three different concentrations (10, 50 and 100 µM). Average speed (µmeters/sec) and total distance were quantified as described in the methods. Hydrogen peroxide (200 mM) was used as a positive control. Asterisks indicate a significant difference (*P* < 0.05) compared to control cells as determined through a one-way ANOVA and Dunnett’s post-hoc test. Data are represented as violin plots including the mean ± quartiles. Figure 4 represent three biological replicates.

### Alkynyl SFN is a more potent and effective antiparasitic compound compared to natural SFN

To determine the binding target of SFN, a novel SFN compound with a single triple bond within the isothiocyanate functional group was synthesized and named alkynyl SFN (A-SFN) (Figure 4A and Supplemental Figure 4). A-SFN was tested against different *Leishmania* species to confirm its effectiveness in reducing viability and inhibiting motility compared to natural SFN. A-SFN was significantly more potent than natural SFN, reducing *Leishmania* viability by greater than 90% against LD and LT at 10 µM (Figure 4B). A-SFN and SFN significantly reduced parasite viability to less than 5% at 50 µM. Supporting a more reactive and effective compound, A-SFN also completely inhibited LT motility after both 3 hours and 6 hours at all concentrations tested (10, 50 and 100 µM) (Figure 4C). These data confirmed that the effects of SFN and A-SFN are comparable, although A-SFN inhibits parasite motility more efficiently and leads to a larger reduction in viability compared to natural SFN.

**Figure 4:**
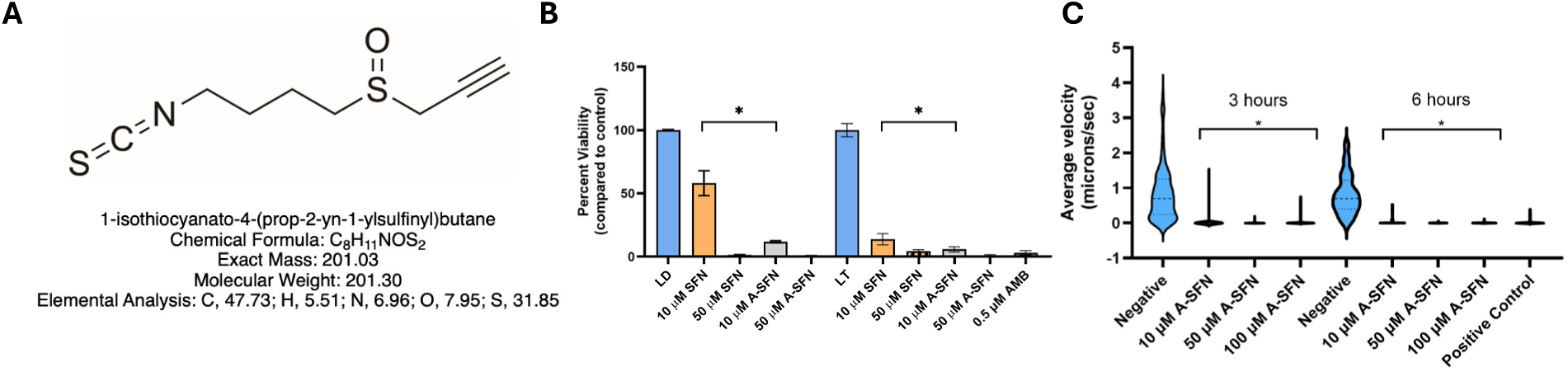
Alkynyl sulforaphane (A-SFN) is a more potent and effective remedial against *Leishmania* compared to natural SFN. **A)** A-SFN chemical structure and mass. **B)** A-SFN decreased promastigote viability against *L. donovani* (LD) [left 5 bars] and *L. tropica* (LT) [right 5 bars] significantly better at 10 µM. 50 µM of either A-SFN or SFN for 48 hours killed greater than 95% of parasites. Viability was quantified as described in Figure 1. Data represent mean ± SD. **C)** LT promastigotes were exposed to 0 (Negative control), 10, 50 or 100 µM of A-SFN for 3 and 6 hours at 27 °C at different concentrations. Average speed (µmeters/sec) was quantified as described in the methods. Asterisks indicate a significant difference (*P* < 0.05) compared to control cells as determined through a one-way ANOVA and Dunnett’s post-hoc test. Data are represented as violin plots including the mean ± quartiles. Figure 4 represent more than three biological replicates.

### KMP-11 sequence conservation and localization via immunofluorescence

Kinetoplastid membrane protein-11 (KMP-11) is a conserved 92 amino acid protein present in all kinetoplastid protozoa [15]. The sequence identity of KMP-11 is high at 92% or greater across all *Leishmania* species compared to 81.5% identity with a KMP-11 expressed in *Trypanosoma brucei* (Figure 5A). Interestingly, 13 out of 92 total residues are lysine residues comprising 14% of the entire KMP-11 protein. KMP-11 is expressed as membrane protein structures on the parasite surface, at the flagellar pocket, and within intracellular vesicles [16]. KMP-11 expression and localization are supported by guided targeting of the KMP-11 gene in *L. mexicana* where two KMP-11 genes (LmxM.34.2220 and LmxM.34.2210) localized expression at the posterior tip and within the entire axoneme [Leishgem.org] [32]. Metacyclic LD promastigotes used for this study strongly expressed KMP-11 at the promastigote surface, within the entire axoneme, and in punctate foci near the flagellar pocket as observed through confocal microcopy (Figure 5B-C).

**Figure 5:**
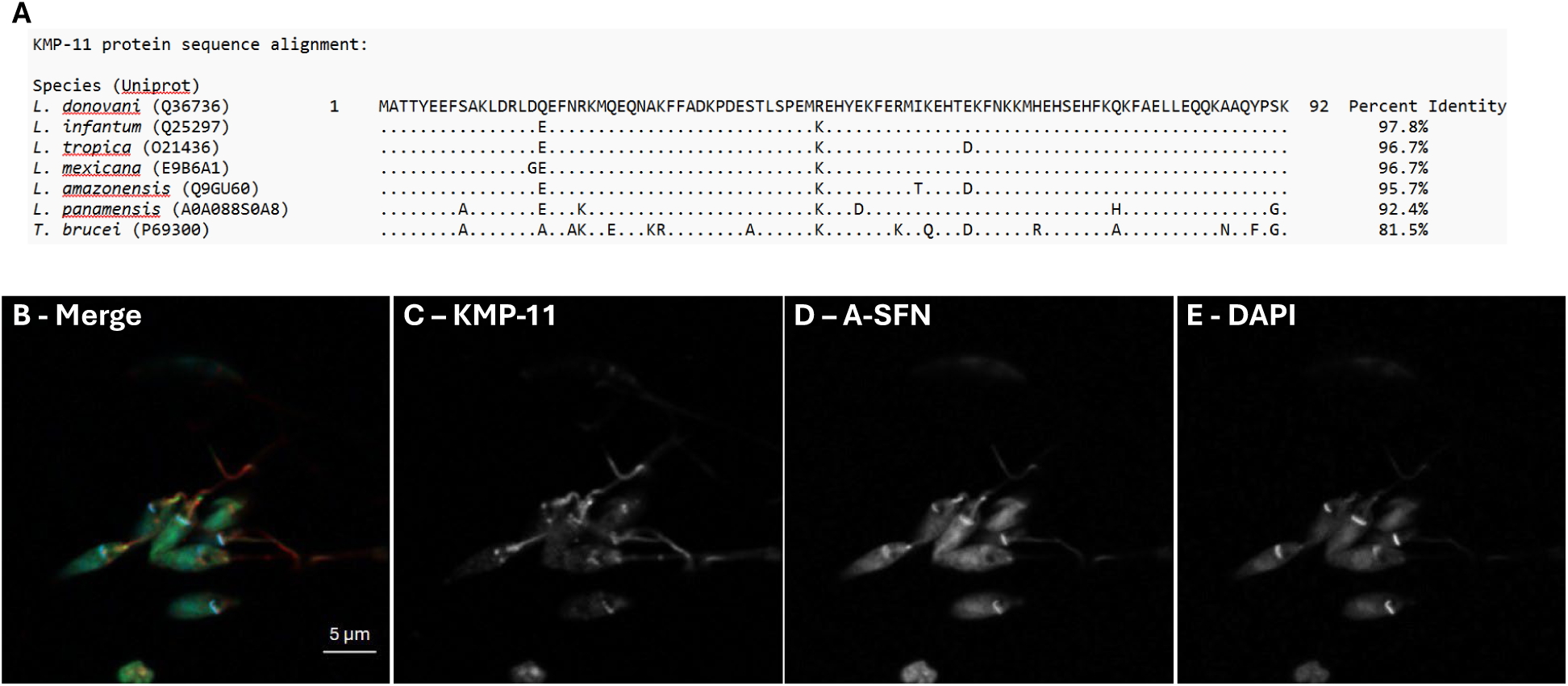
KMP-11 is conserved across *Leishmania* and is not bound by A-SFN. **(A)** The amino acid sequence of KMP-11 is identified by their accession numbers from *L. donovani* (LD)*, L. tropica* (LT)*, L. infantum* (LI)*, L. tropica* (LT), *L. mexicana* (LM)*, L. amazonensis* (LA), *L. panamnesis* (LP) or from *Trypanosoma brucei* (TB) with single amino acid substitutions indicated from the LD KMP-11 sequence. **(B-E)** Localization of KMP-11 (C) and A-SFN (D) in metacyclic promastigotes was identified through CuAAC chemistry (SFN labelled green) and immunofluorescence (KMP-11 labelled red). KMP-11 expression was along the axoneme, at the posterior tip and strong expression at the flagellar pocket. DAPI was used to stain the kinetoplast (E).

### SFN covalently binds *Leishmania*

Synthesis of A-SFN, which contains a single triple bond on the opposing end from the reactive isothiocyanate group, enabled click chemistry reactions to be conducted to perform site-specific protein labeling on the surface of living parasites to identify possible binding targets (Uttamapinant et al., 2012). click reactions were also used to confirm binding through a modified *in vitro* binding assay [30]. Guided by the motility data that indicates SFN inhibits the parasite within the first three hours at 100 µM, LM promastigotes were exposed to 100 µM A-SFN for 3 hours, fixed, and labeled through CuAAC reactions to identify the localization of A-SFN, which was covalently bound to a structural analog of Alexa Fluor™488 (green fluorescence). Simultaneously, the cellular localization of KMP-11 was confirmed through immunofluorescence utilizing Alexa Fluor™594 (red fluorescence) and used to assess the degree of colocalization with A-SFN. KMP-11 localized at the promastigote surface, within the axoneme, and in punctate foci near the flagellar pocket (Figure 5 C). A-SFN localized mainly within the cytoplasm, but was also highly expressed within the flagellar pocket (Figure 5D). Analysis of the Mander’s coefficient analysis revealed that A-SFN colocalized with KMP-11 of 20% (Figure 5B). Although the immunofluorescence click experiment provided initial evidence that SFN binds the promastigotes, the results were not conclusive that KMP-11 was the true target and did not rule out the interaction of other binding partners. Therefore, A-SFN was incubated with recombinant KMP-11 through an *in vitro* binding experiment utilizing click chemistry linked to a biotin-azide that was immunoprecipitated with avidin beads. However, the binding assay did not confirm binding of KMP-11 and SFN.

### SFN activates the NRF2 pathway in macrophages post-infection inducing autophagic changes

*Leishmania* parasites that cause VL require NRF2 activation post-infection to combat lipid peroxidation and ferroptosis [33]. Therefore, SFN treatment early in infection may modulate the NFR2 pathway and negatively affect the parasite lifecycle by altering antioxidant or other genes important for amastigote differentiation. To begin to elucidate these alterations, global RNA transcriptomic analysis identified several basal and inducible host NRF2-dependent genes that were elevated in expression in infected cells treated with SFN compared to infection alone after 24 hours (Table 1). SFN has been previously shown to increase lysosomal acidification by activating TFEB, a master regulator of autophagic and lysosomal functions, which leads to enhanced expression of genes required for autophagosome and lysosome biogenesis [34]. Indeed, in LD-infected cells treatment with SFN significantly increased the expression of autophagic genes including SQSTM1 (Fold Change (FC) increase of 1.6), ULK1 (FC increase of 1.4), and MAP1LC3B (FC increase of 1.3) as previously described [34]. Uninfected control cells treated with SFN had similar increases in expression (SQSTM1 FC = 1.7, ULK1 FC = 1.7 and MAP1LC3B FC = 1.6) supporting our results. Moreover, enhanced protein expression of SQSTM1 in LD-infected cells was confirmed after 48 hours of SFN treatment (Supplemental Figure 5). Therefore, SFN treatment given after LD infection activates the canonical type I antioxidant and detoxification pathway as well as a type II autophagic and lysosomal pathway after *Leishmania* infection.

**Table 1:**
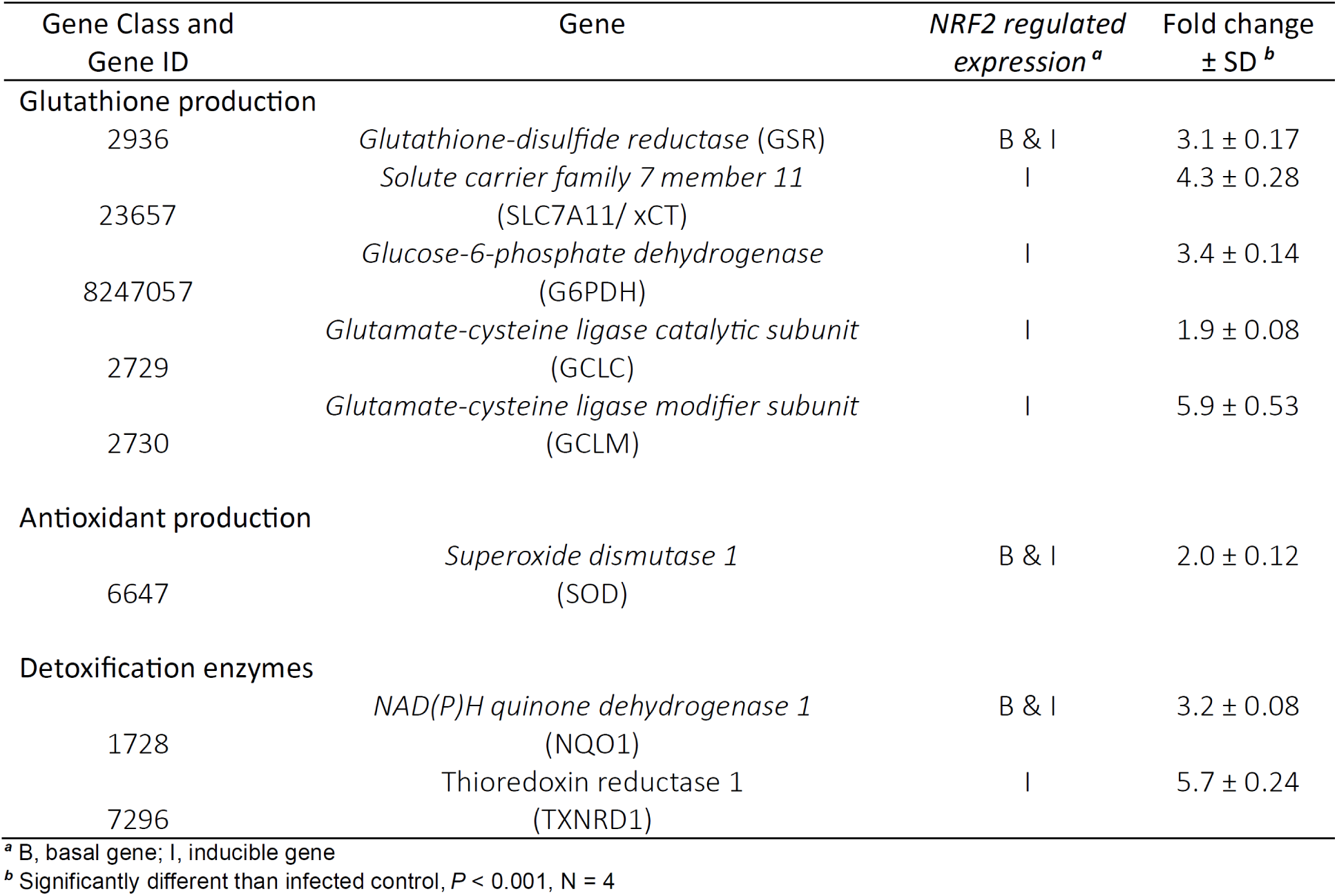
Sustained NRF2 upregulated gene expression in THP1 cells during *Leishmoina donovoni* infection in the presence of 10 μM SFN after 24 hours.

To confirm our transcriptomics data mechanistically, autophagic flux was assessed early during L*eishmania* infection in the presence of SFN. THP-1 cells were stained with a small, pH-sensitive marker (DALGreen) prior to infection (5 hours) then treated with or without SFN for an additional 18 hours. DALGreen is a small cell-permeable fluorescence probe that detects autolysosomes and increases fluorescence at acidic pHs [35]. LI infection led to a significant decrease in the median DALGreen fluorescence (Figure 6C) as well as in the number of DALGreen^+^ cells after 24 hours (Figure 6D), indicating that LI alters autophagic flux and inhibit autolysosome (AL) formation early post-infection as previously described (Thomas et al., 2018).

**Figure 6:**
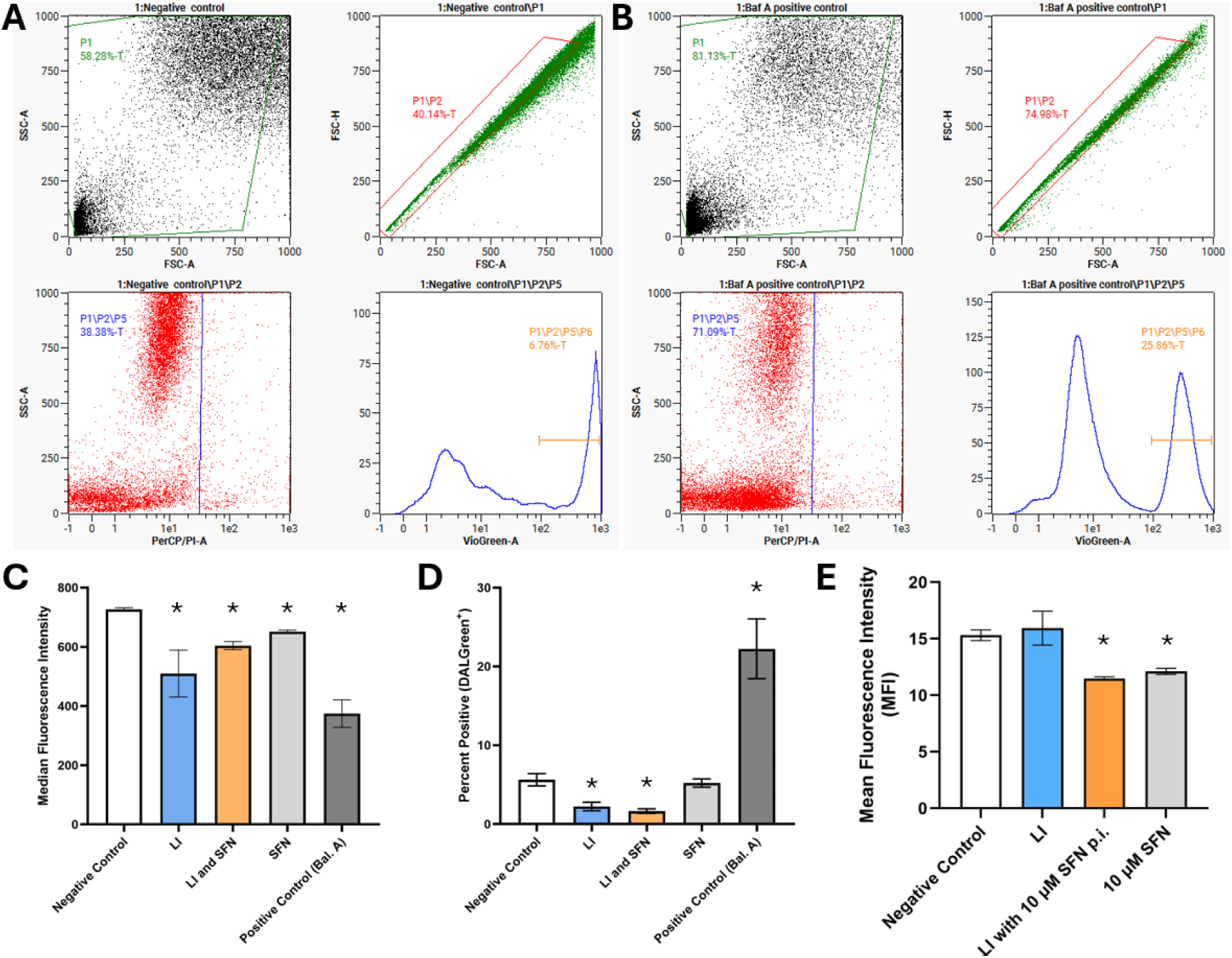
Sulforaphane alters autophagic flux post-infection. THP1 macrophages were stained with DALGreen, infected with LI then exposed to SFN (10 μM) for 18 hours. DALGreen fluorescence was quantified in viable, single cells in negative control cells **(A)** or in cells treated with Bafilomycin A (100 nM) **(B)** 24 hours post-infection. **C)** Median fluorescence and **D)** the percentage of DALGreen positive cell were quantified as described in the methods. THP1 macrophages were also infected with LI then exposed to SFN (10 μM) for 15 hours then treated with 50 μg/ml FITC-dextran for 3 hours. FITC fluorescence was also quantified in viable, single cells, and the **E)** Mean fluorescence was quantified as described in the methods. Asterisks indicate a significant difference (*P* < 0.05) compared to control cells as determined through a one-way ANOVA and Dunnett’s post-hoc test. Data are represented as mean or median ± SD. Figure 6 represent three biological replicates.

SFN treatment post-infection significantly increased DALGreen fluorescence compared to LI infection alone, but did not alter the percentage of DALGreen^+^ cells. These data indicate an equivalent number of ALs 24 hours post-infection with more acidic lumen content induced by SFN treatment. MΦ’s treated with SFN alone in the absence of infection had higher levels of DALGreen fluorescence compared to infected cells after 18 hours indicating that SFN by itself alters autolysosome acidification. Bafilomycin A (BAL), an established inhibitor of v-ATPase activity, inhibits lysosomal acidification at early time points and inhibits the fusion of autophagosomes (APs) and lysosomes during later time points [36–38]. BAL treatment (100 nM for 18 hours) led to a significant decrease in DALGreen fluorescence correlated with increased pH in ALs (Figure 6C and D). Moreover, the percentage of DALGreen^+^ cells was significantly increased with BAL due to the inhibition of fusion (Figure 6E). These results confirm that VL parasites, such as LI, inhibit autophagy early during infection and that SFN treatment increases lysosomal pH post-infection.

SFN increasing lysosomal acidification was also confirmed post-infection by assessing lysosomal acidification using a pH sensitive dextran. Fluorescein isothiocyanate (FITC)-labeled dextran is a molecule used to assess phagocytotic activity and whose fluorescence diminishes with lysosomal acidification [39]. THP-1 cells were infected with LI for 5 hours, treated with or without SFN for 12 hours, then treated with FITC-labeled dextran (size: 70 kD, 60 Å) for 3 hours (Figure 6E). We observed by flow cytometry that FITC fluorescence diminished in an SFN-dependent manner irrespective of infection, confirming that SFN enhances lysosomal acidification.

## 7. DISCUSSION and CONCLUSIONS

Infection of *Leishmania donovani* or *L. infantum* leads to the development of visceral leishmaniasis (VL) in humans that is often fatal if untreated. VL is a NTD that is associated with poverty, malnutrition, and rural populations who typically lack appropriate healthcare options (Aronson et al., 2026; World Health Organization, 2023). The current drugs used to treat VL are highly toxic, expensive and are ineffective in certain regions due to resistant parasites, thereby limiting an already small group of antileishmanial therapies [5]. Leishmaniasis is typically diagnosed in Africa, Asia and the Middle East, however, leishmaniasis is now endemic in the US with the prevalence of leishmaniasis increasing in the US [8–11,41]. *Leishmania* infections are predicted to increase worldwide with endemic US infections expected to double by 2080 driven by climate change [42]. Unfirtunately, the number of new chemical entities that are suitable for clinical development for the treatment of VL remains low [43]. For these reasons, new treatments against VL and other forms of leishmaniasis are urgently needed worldwide.

Our approach, based on the One Health paradigm [6,7], initially identified SFN as an effective anti-leishmanial compound that kills LD and LI promastigotes through an unknown mechanism that was independent of host or parasite ROS [23]. SFN is a well-known ITC, which is a class of anticarcinogenic compounds first described as potent inducers of phase II enzymes [22]. ITCs, like SFN, are also reactive electrophiles due to the presence of the isothiocyanate group (N=C=S) that can covalently modify cysteine and lysine residues through addition reactions [44,45]. ITCs predominantly target lysine and cysteine R groups, although in some cases amino acids such as tyrosine, threonine and serine may also be modified [46]. The isothiocyanate functional group is known to react irreversibly with thiol side chains of cysteines to form dithiocarbamates and with the primary amine of lysine side chains to form thioureas [46]. However, this reaction requires a pH 9.0–11.0 for optimal conjugation [46]. Cysteine side chains are better nucleophilic targets, but the binding of ITCs to cysteine are unstable and have been shown to be reversible [47]. Numerous examples of covalent modifications of proteins with various isothiocyanates have been reported in the literature. For example, SFN has been shown to covalently bind acyl-protein thioesterase 2 (APT2) through Cys56, while phenethyl isothiocyanate (PEITC) binds MEKK1, an upstream regulator of the SAPK/JNK signal transduction pathway, through Cys1238 [48,49].

Covalent modifications induced by SFN targeting cysteine residues, specifically Cys151 on KEAP1, have been shown to cause dissociation of KEAP1 from its counterpart NRF2. NRF2 is a well-known transcription factor that induces the expression of antioxidant and detoxification (phase II) enzymes once liberated from its cellular inhibitor (Dinkova-Kostova et al., 2002; Kensler et al., 2007). Recent data from our lab indicates that in the *in vitro* MΦ model used in this infection study, SFN is a broad cellular activator with low toxicity at 10 µM after 48 hours [24]. Moreover, SFN activates a broad transcriptional signature, upregulating more than 600 genes compared to less than 200 genes activated by more specific, noncovalent PPIs [24]. Together SFN and PPI inhibtors upregulated a core group of approximately 50 known Nrf2-dependent genes in MΦ’s. These data indicate that SFN can participate in both NRF2-dependent and NRF2-independent mechanisms of action within infected cells. This complexity and the contribution of NRF2-dependent (KEAP1 inhibition) and NRF2-independent (covalently modification of cysteine and lysine residues) mechanisms need to be further examined to define the complete mechanism of SFN parasitic killing.

During successful *Leishmania* infections, the parasite subverts host proteins to modify the intracellular cellular environment to permit parasite replication [50]. For example, NRF2 has been recently shown to be a critical transcription factor that protects LI parasites from lipid peroxidation and death through ferroptosis by inducing an NRF2-dependent antioxidant response in primary murine macrophages [33,51,52]. Therefore, it follows that therapeutic approaches to modify the activation status of NRF2 may counteract ferroptosis resistance driven by antioxidant defenses and restore host antipathogen functions [53]. To further complicate this essential cellular stress pathway, many NRF2-dependent genes are linked with macroautophagy [54,55], which is essential and temporally controlled during VL infections [56]. For example, infection of human MΦ’s with LD is known to inhibit canonical autophagy initially (< 24 hours post-infection), then activates noncanonical autophagy later. Interestingly, many NRF2-dependent antioxidant, detoxification and glutathione synthesis enzymes have been previously hypothesized to inhibit and not enhance VL infections [57]. Altogether these contradictory results at first seem counterintuitive, the complexity of the NRF2 pathway as a master regulator of the cellular stress reveals that modification of this keystone pathway is not straight forward [58]. Therefore, we hypothesize that altering the timing or magnitude of this pathway post-infection with SFN will negatively affect parasite differentiation and replication.

Understanding SFN’s broad intracellular effects in host macrophages informed our investigation into possible protein targets of *Leishmania* focused on essential parasite proteins enriched in lysine and cysteine residues. KMP-11 was identified as an initial target. KMP-11 has a high percentage (∼15%) of lysine residues that participate in phase separation [59]. Moreover, KMP-11 is essential in regulating the overall lipid bilayer of the parasite membrane (Jardin 1995). Most recently, KMP-11 was elegantly shown to regulate cholesterol transport from the host to the parasite and acted as an essential protein that interacts with the host 1) to create a bridge to transport host cholesterol to the parasite, and 2) is essential for LD to attach and invade macrophages [21]. Interestingly, KMP-11 was predicted *in silico* to participate in hydrogen bonding with KMP-11 proteins across many *Leishmania* species possibly through its lysine residues [60]. Therefore, SFN binding to KMP-11 was directly tested through immunofluorescence and an *in vitro* binding assay.

Our data indicate that SFN was able to inhibit *Leishmania* promastigote motility after 6 hours in a dose-dependent manner leading to decreased viability after 48 hours. Moreover, our newly synthesized A-SFN with a triple bond, alkyne group, was more potent than natural SFN in all endpoints quantified. Previously several different synthetic derivatives of sulforaphane have been synthesized and the structure activity relationships between the sulfoxide side chain and cytotoxicity has been reported [28]. Increasing the length of the alkyl chain from methyl in sulforaphane to a longer, more nonpolar side chains led to increased cytotoxicity. While the alkynyl side group in our SFN-A derivative was not specifically studied, the increased activity of SFN-A compared to SFN fits with prior data by Ren and coworkers showing a link between increased side chain lipophilicity and increased biological activity [28]. Although A-SFN was more potent, KMP-11 was not a confirmed binding target *in vitro*. Therefore, future experiments will utilize A-SFN and click chemistry through immunoprecipitation and proteomic analysis.

## 8. ACKNOWLEDGEMENTS

We would like to acknowledge the effort and time of Cora Tunnel, Everly Ivener, Haley Dee and Tanisha Garcia who were undergraduate Biology and Chemistry majors at Fort Lewis College that participated in this NTD project. The following reagents were obtained through the BEI Resources, NIAID, NIH: *Leishmania mexicana,* Strain MNYC/BZ/62/M379 (+luc), NR-51210, *Leishmania amazonensis*, Strain LV78 (+bla), NR-49247, and *Leishmania panamensis*, Strain PSC-1 (MHOM/PA/94/PSC 1), NR-50162.

## 9. FINANCIAL SUPPORT

Research reported in this publication was supported by the National Institute of Allergy and Infectious Diseases of the National Institutes of Health under Award Number (1R16AI189394-01) (DB). This work was supported by AIM center cores funded by NIH grant P20GM121176. Research was also supported by the Beckman Scholars Program through the Arnold and Mabel Beckman Foundation (EK). The Fort Lewis College Undergraduate research fund supported reagents used by AH and KT. This research was supported by a National Science Foundation – Major Research Instrumentation award (2017945). The authors wish to thank the Analytical Resources Core (RRID: SCR_021758) at Colorado State University for instrument access, training and assistance with sample analysis (KM).

## Notes

### Competing Interest Statement

The authors have declared no competing interest.

