## Supplemental Figures for "Sulforaphane targets *Leishmania* promastigotes to inhibit parasite motility and alters host autophagy to suppress parasite replication in macrophages"

### Supplemental Data:

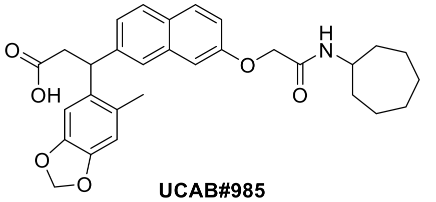

UCAB#985

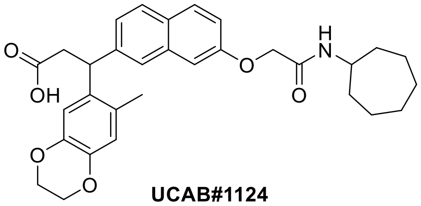

UCAB#1124

**Supplemental Figure 1.** Chemical structures of the two previously published non-covalent Keap1-Nrf2 inhibitors tested in this study.

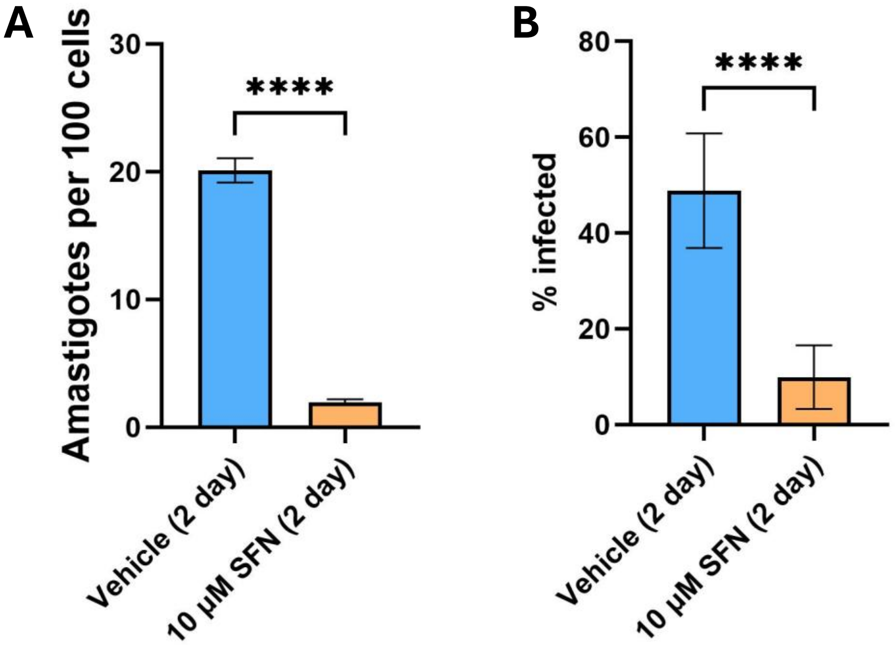


**Supplemental Figure 2:** Sulforaphane is an effective therapeutic in macrophages stably infected with *Leishmania amazonensis* (LA). SFN treatment decreases the number of intracellular amastigotes in macrophages **(A)** and the percent of infected cells **(B)** after 24-hour infection and 48 hours of exposure of SFN*.* Asterisks indicate a significant difference ($P < 0.05$) compared to control cells as determined through a one-way ANOVA and Dunnett's post-hoc test. Data represent mean ± SD and one biological replicate.

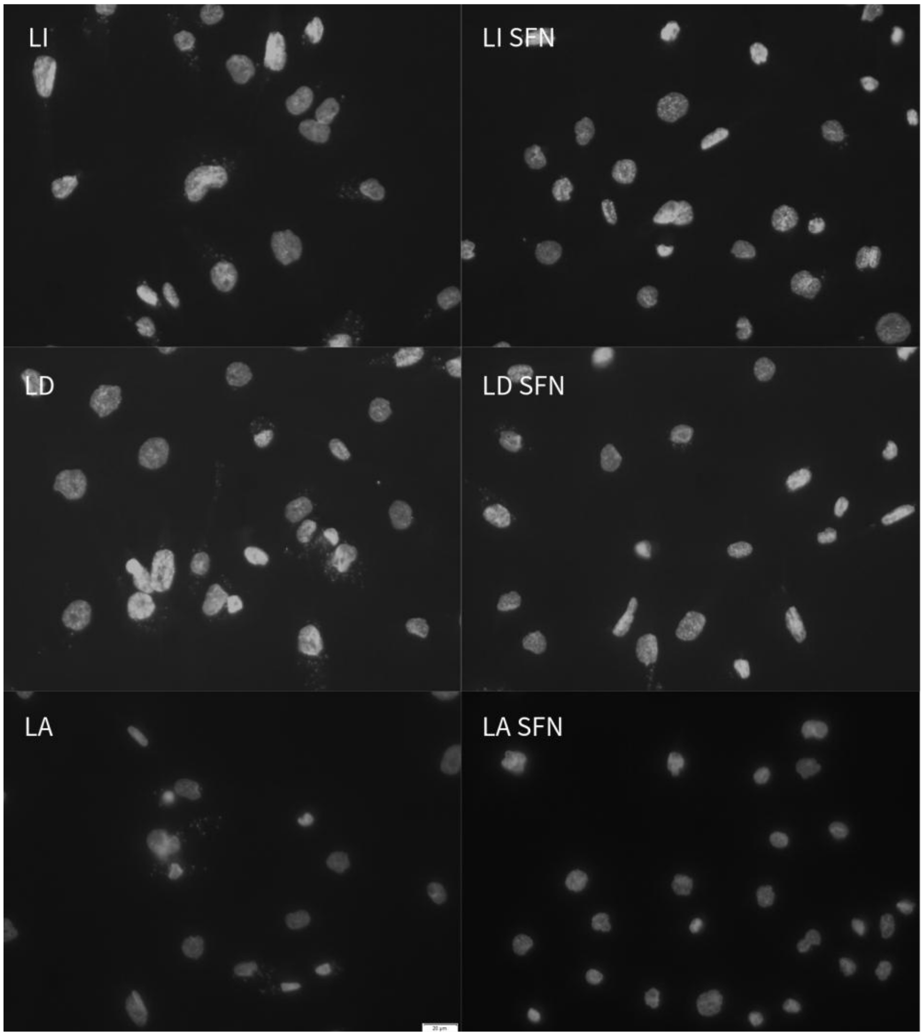


**Supplemental Figure 3:** Representative immunofluorescence images of macrophages (large nuclei) stably infected with *Leishmania donovani* (LD)*, infantum* (LI)*, and amazonensis* (LA) (small punctate foci)**.** SFN treatment decreases the number of intracellular amastigotes in macrophages after 24-hour infection and 48 hours of exposure of SFN*.*

**Supplemental NMR Methods:**

All chemicals were purchased from Sigma-Aldrich and used without further purification. All solvents were degassed with nitrogen and passed through activated molecular sieves prior to use. Reactions involving air or moisture sensitive reagents or intermediates were performed under an inert atmosphere of nitrogen in glassware that had been oven or flame dried. The $^{1}H$ and $^{13}C$ NMR spectra were obtained on a Bruker Avance NEO-400 spectrometer, operating at 400 and 100 MHz respectively. Unless indicated otherwise, all spectra were run as solutions in $CDCl_3$. The $^{1}H$ NMR chemical shifts are reported in parts per million (ppm) downfield from tetramethylsilane (TMS) and are, in all cases, referenced to the residual protio-solvent present (d 7.24 for $CHCl_3$). The $^{13}C$ NMR chemical shifts are reported in ppm relative to the center line of the multiplet for deuterium solvent peaks (d 77.0 (t) for $CDCl_3$).

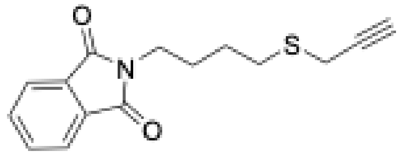


**2-(4-(prop-2-yn-1-ylthio)butyl)isoindoline-1,3-dione**. NaH (60% dispersion in mineral oil, 0.096 g, 2.4 mmol) was added in one portion to a solution of 2-(4-mercaptobutyl)isoindoline-1,3-dione (0.475 g, 2.0 mmol) in THF (10 mL) at 0 °C and the solution was stirred for 30 min. A solution of propargyl bromide (80 wt. % in toluene, 0.356 g, 2.4 mmol) was then added by syringe and the solution was stirred at rt for 22 h. The reaction was quenched with water (10 mL) and extracted with EtOAc (3 x 15 mL). The combined organic layers were dried ($MgSO_4$), filtered, and concentrated under reduced pressure. The residue was purified by flash chromatography eluting with EtOAc/hexane (1:9) to give 0.375 g (68%) as a pale-yellow oil; $^{1}H$ NMR (400 MHz) 7.83 (dd, $J$ = 5.2, 2.8 Hz, 2H), 7.20 (dd, $J$ = 5.2, 2.8 Hz, 2H), 3.72 (t, $J$ = 6.8 Hz, 2H), 3.23 (d, $J$ = 2.8 Hz, 2H), 2.73 (t, $J$ = 7.2 Hz, 2H), 2.22 (t, $J$ = 2.4 Hz, 1H), 1.85-1.78 (m, 2H), 1.72-1.64 (m, 2H); $^{13}C$ NMR (100 MHz) 168.4, 133.9, 132.0, 123.2, 80.0, 71.0, 37.4, 31.0, 27.7, 26.2, 19.1.

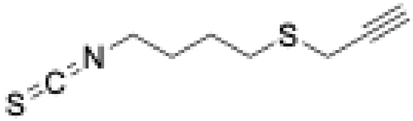

**(4-isothiocyanatobutyl)(prop-2-yn-1-yl)sulfane**. Hydrazine hydrate (0.172 g, 0.166 mL, 3.43 mmol) was added to a solution of 2-(4-(prop-2-yn-1-ylthio)butyl)isoindoline-1,3-dione (0.375 g, 1.37 mmol) in MeOH (9 mL) at rt, and the reaction was stirred for 3 h. The reaction was concentrated under reduced pressure and a 1M solution of NaOH (15 mL) was added. The solution was extracted with EtOAc (4 x 15 mL), and the combined organic layers were dried ($MgSO_4$), filtered, and concentrated under reduced pressure to give the primary amine (0.178 g, 90%) as a colorless oil. The oil was dissolved in THF (9 mL) and cooled to 0 °C. $Et_3N$ (0.446 g, 4.42 mmol) and $CS_2$ (0.115 g, 1.52 mmol) were added. The reaction was warmed to rt and stirred 2 h. The reaction was cooled to 0 °C and MsCl (0.174 g, 1.52 mmol) was added. The reaction was warmed to rt and was stirred for 30 min. $CH_2Cl_2$ (15 mL) was added, and the mixture was washed with 1N HCl (3 x 5 mL), water (5 mL), and brine (5 mL). The organic layer was dried ($MgSO_4$), filtered, and concentrated under reduced pressure. The residue was purified by flash chromatography eluting with EtOAc/hexane (1:10) to give 0.168 g (73%) as a pale-yellow oil which solidified on standing; $^1$H NMR (400 MHz) 3.59 (t, *J* = 6.0 Hz, 2H), 3.28 (d, *J* = 2.4 Hz, 2H), 2.76 (t, *J* = 6.8 Hz, 2H), 2.28 (t, *J* = 2.8 Hz, 1H), 1.90-1.76 (m, 4H)Hz; $^{13}$C NMR (100 MHz) 130.4, 79.8, 71.2, 44.7, 30.7, 28.9, 25.8, 19.2.

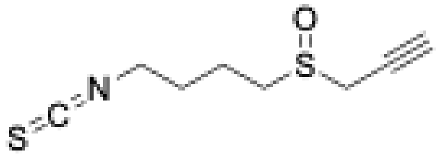

**1-isothiocyanato-4-(prop-2-yn-1-ylsulfinyl)butane**. A solution of MCPBA (0.093 g, 0.54 mmol) in $CH_2Cl_2$ (1.6 mL) was added dropwise over 5 min to a solution of (4-isothiocyanatobutyl)(prop-2-yn-1-yl)sulfane (0.100 g, 0.54 mmol) in $CH_2Cl_2$ (4 mL) at -10 °C, and the reaction stirred at -10 °C for 1 h. A saturated solution of $NaHCO_3$ (15 mL) was added and the mixture was extracted with $CH_2Cl_2$ (2 x 15 mL). The combined organic layers were washed with saturated $NaHCO_3$ (30 mL), brine (30 mL), dried ($MgSO_4$), filtered, and concentrated under reduced pressure. The residue was purified by flash chromatography eluting with MeOH/$CH_2Cl_2$ (1:20) to give 0.084 g (77%) as a colorless oil; $^1$H NMR (400 MHz) 3.63 (t, *J* = 6.4 Hz, 2H), 3.62 (d, *J* = 2.4 Hz, 2H), 3.02-2.89 (m, 2H), 2.49 (t, *J* = 2.4 Hz, 1H), 2.06-1.88 (m, 4H); $^{13}$C NMR (100 MHz) 131.2, 76.7, 72.3, 50.4, 44.6, 42.1, 29.0, 19.8.

**Supplemental NMR Figure 4:**

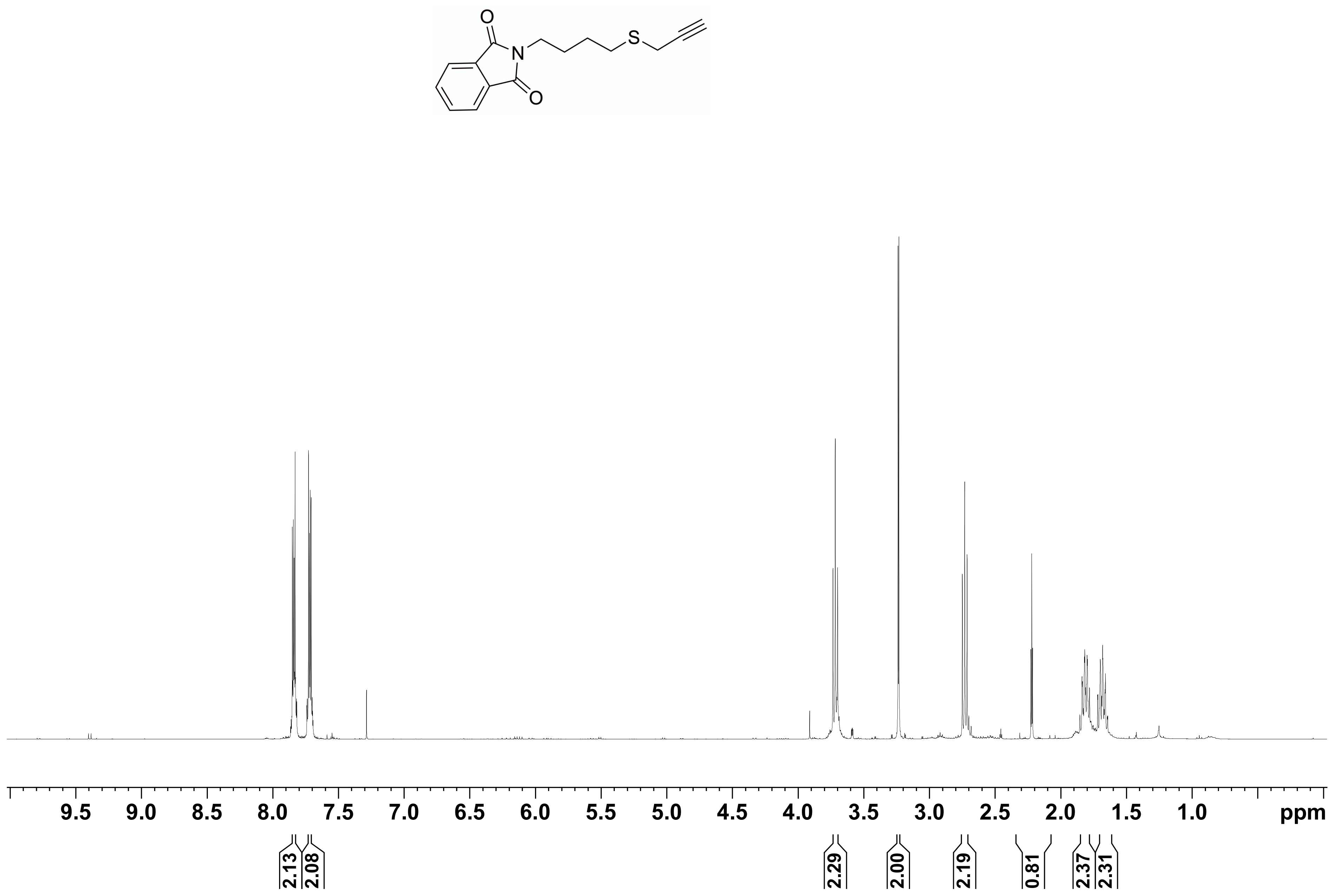

O
N
S
O
9.5
9.0
8.5
8.0
7.5
7.0
6.5
6.0
5.5
5.0
4.5
4.0
3.5
3.0
2.5
2.0
1.5
1.0
ppm
2.13
2.08
2.29
2.00
2.19
0.81
2.37
2.31

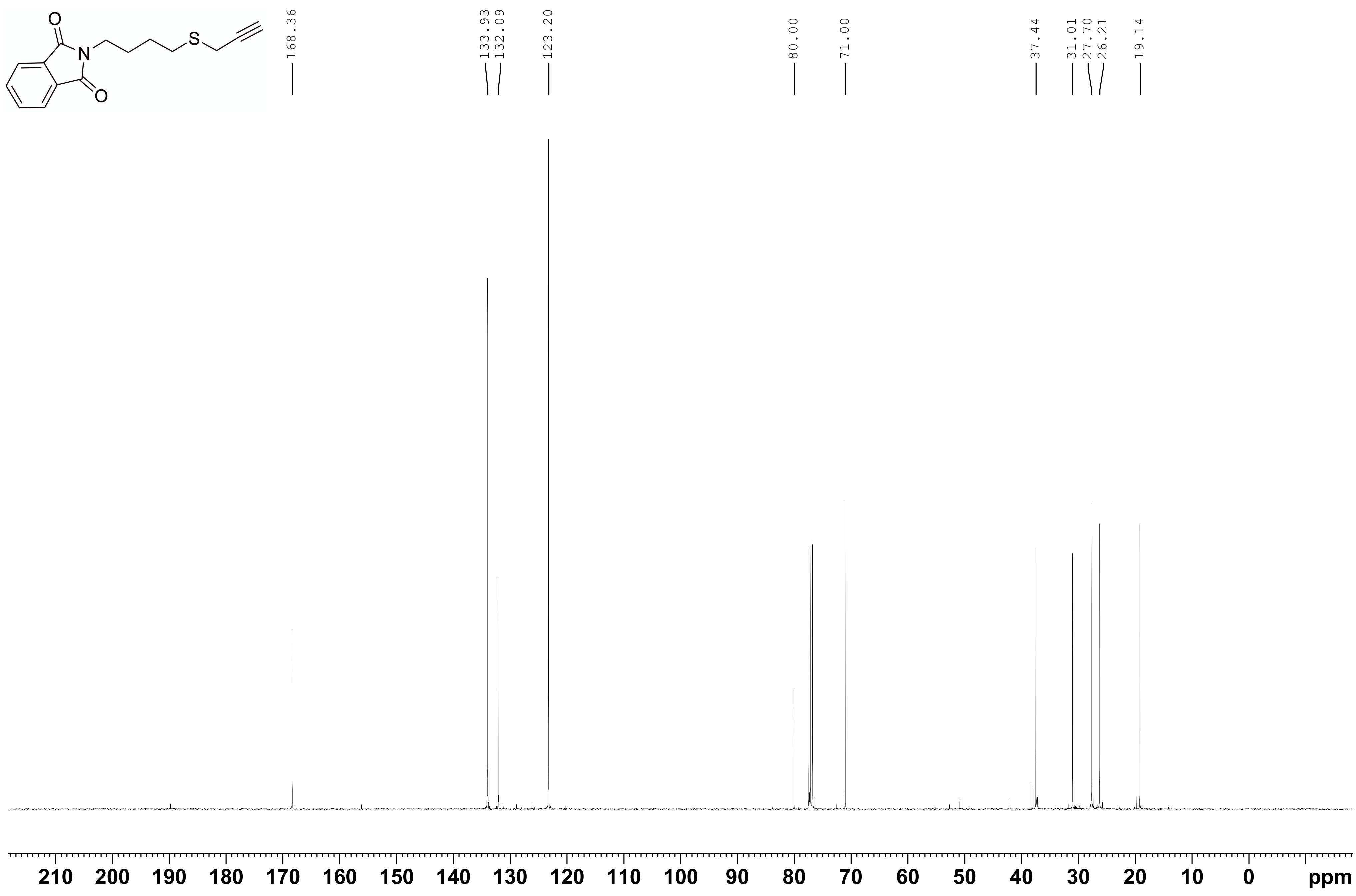

S
N
O
O
168.36
133.93
132.09
123.20
80.00
71.00
37.44
31.01
27.70
26.21
19.14
210 200 190 180 170 160 150 140 130 120 110 100 90 80 70 60 50 40 30 20 10 0
ppm

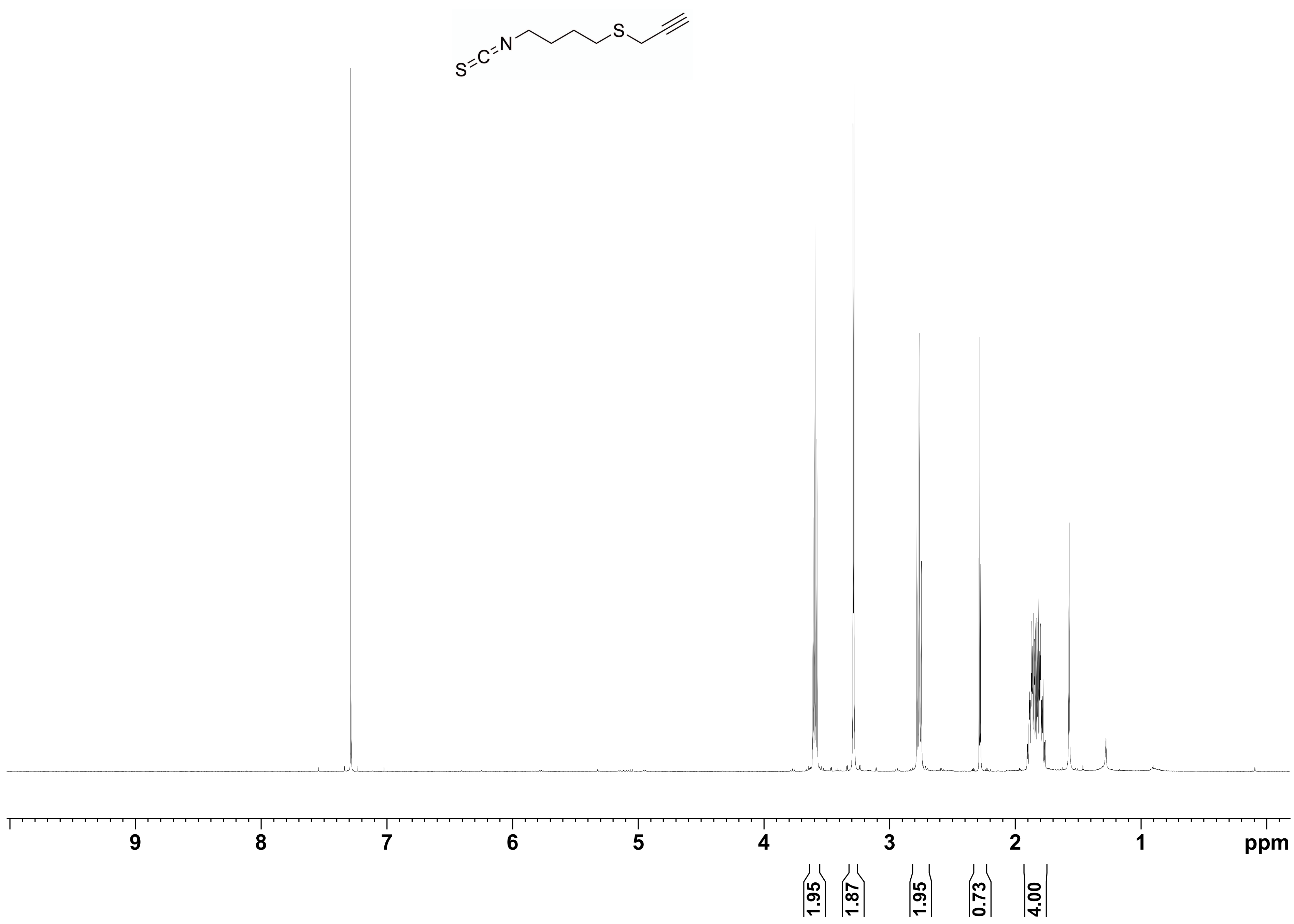
S=C=N
S
9
8
7
6
5
4
3
2
1
ppm
1.95
1.87
1.95
0.73
4.00

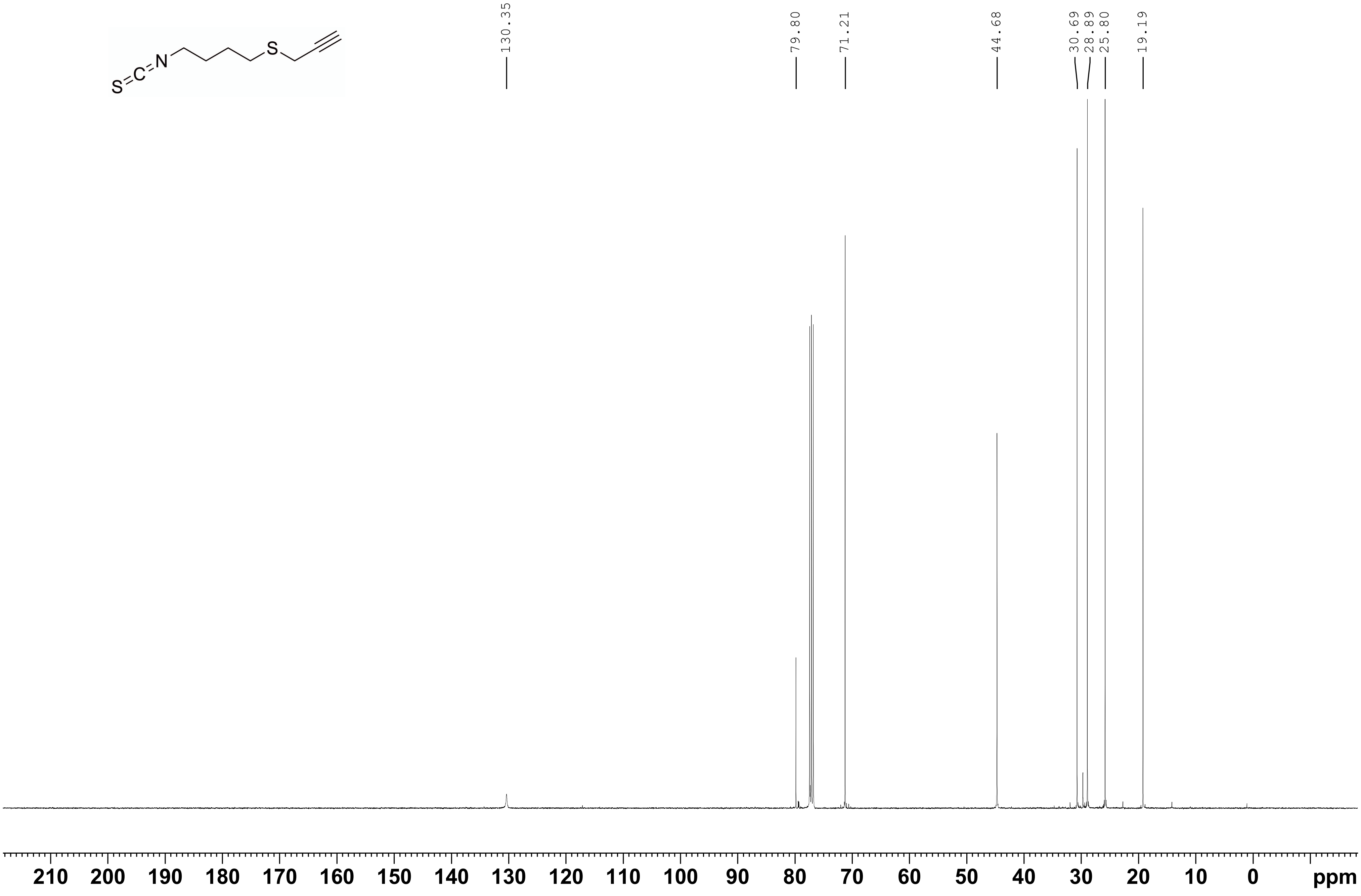

S=C=N
S
130.35
79.80
71.21
44.68
30.69
28.89
25.80
19.19
210
200
190
180
170
160
150
140
130
120
110
100
90
80
70
60
50
40
30
20
10
0
ppm

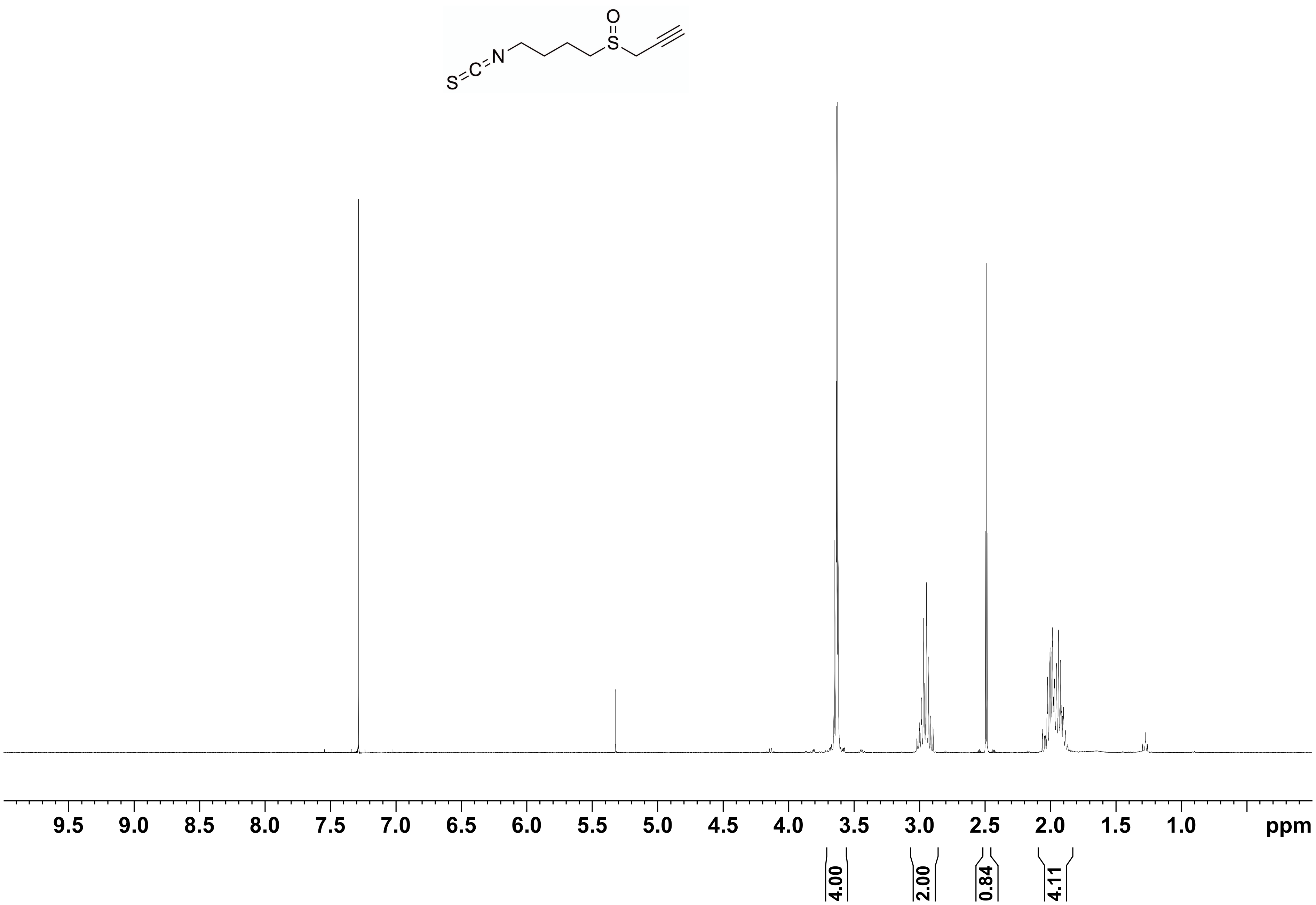

S=C=N
S
O
9.5
9.0
8.5
8.0
7.5
7.0
6.5
6.0
5.5
5.0
4.5
4.0
3.5
3.0
2.5
2.0
1.5
1.0
ppm
4.00
2.00
0.84
4.11

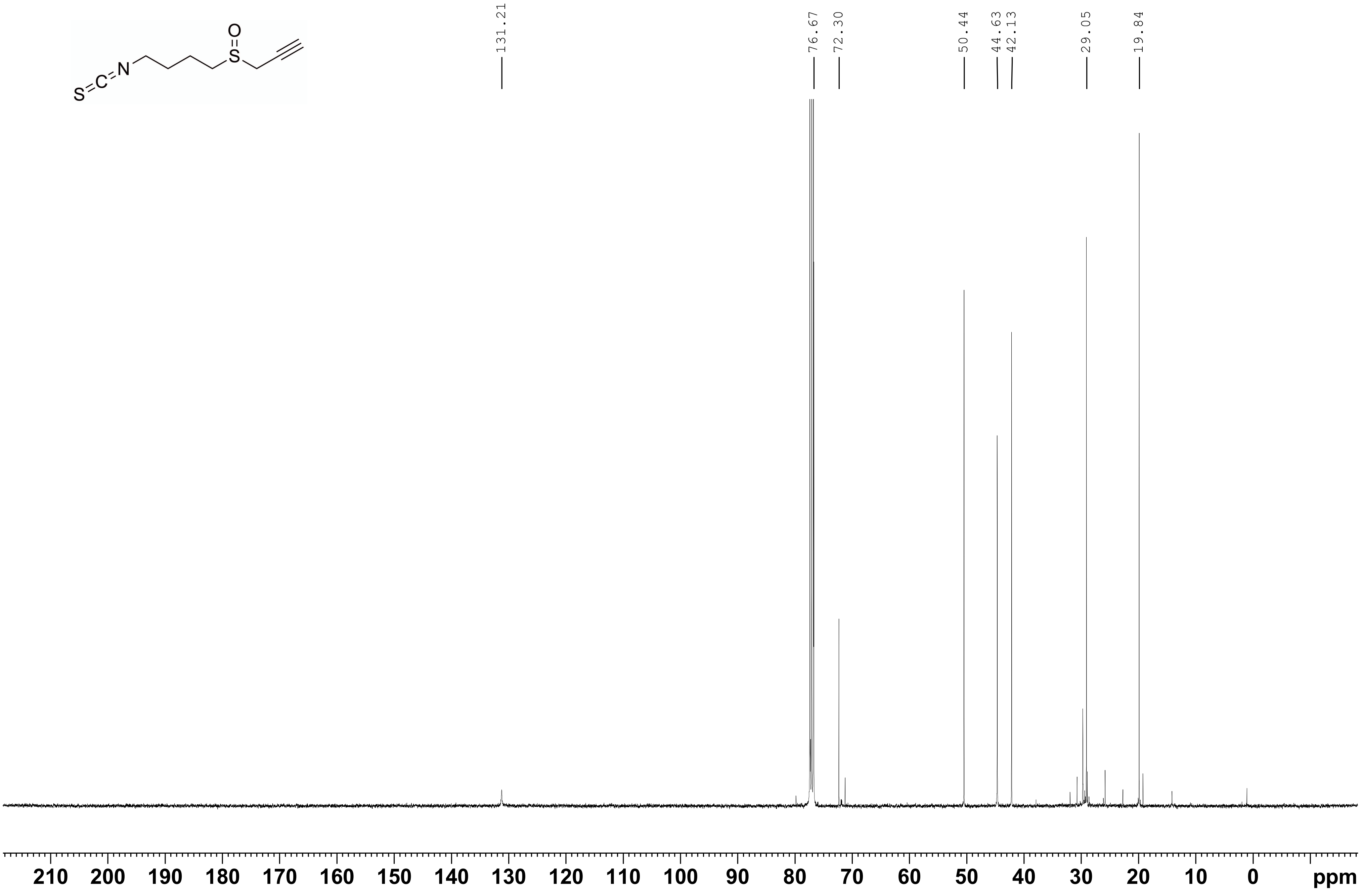
131.21
76.67
72.30
50.44
44.63
42.13
29.05
19.84
210 200 190 180 170 160 150 140 130 120 110 100 90 80 70 60 50 40 30 20 10 0
ppm

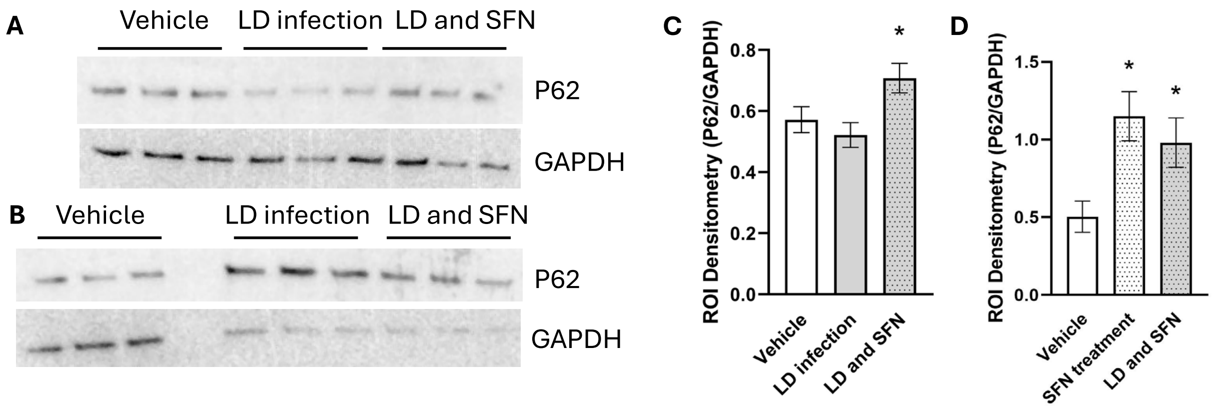


**Supplemental Figure 5:** Macrophages infected with *Leishmania donavani* (LD) and treated with SFN for 48 hours upregulated P62 expression. **(A and B)** Representative western blots of THP-1 cells treated with SFN. **(C and D)** Protein quantification of western blots in A and B. Asterisks indicate a significant difference ($P < 0.05$) compared to control cells as determined through a one-way ANOVA and Dunnett's post-hoc test. Data represent mean ± SD and one biological replicate.
